# Low-Cost 3D Printed Optics for Super-Resolution Multifocal Structured Illumination Microscopy

**DOI:** 10.1101/2025.10.09.680962

**Authors:** Jay Christopher, Liam M. Rooney, Charlie Butterworth, Gail Mcconnell, Ralf Bauer

## Abstract

We present a low-cost 3D printing method of fabricating optical quality lenslet arrays for integration in a multifocal structured illumination microscope (mSIM), achieving fluorescence imaging below the optical diffraction limit. We detail the design and manufacturing processes to produce high-quality 3D printed optics, showing their comparable surface roughness of 30 ± 2.5 nm for the 3D printed elements compared to 37 ± 1.4 nm for commercial glass optics. A 3D printed lenslet array with a ‘honeycomb’ geometry and 1.2 mm lenslet diameter was compared to a high-end glass commercial lenslet array with 250 µm lenslet diameter and a lower cost commercial lenslet array with a 1.2 mm by 1.6 mm lenslet footprint. The imaging performance of the different optics was benchmarked using a custom mSIM setup by quantifying the beam profile homogeneity and the experimental lateral resolution. The mSIM setup incorporating the different microlens arrays was tested using a commercial bovine pulmonary artery endothelial cell specimen, highlighting an achievable resolution enhancement from 229 nm ± 11 nm with widefield illumination to 137 ± 11 nm using the high-end commercial microlens array and 134 nm ± 9 nm using the 3D printed honeycomb lenslet array. Advantages of improved background rejection through the custom lenslet geometry are discussed, highlighting the super-resolution microscope performance achievable through custom low-cost 3D printed optics.

## 1. Introduction

Biological imaging research remains fundamental for enabling new advances in both life sciences and healthcare. The latest developments of novel microscopy approaches are pursued in tandem with initiatives to make low-cost and open-source microscopy developments more widely available [1–3], and recently the potential to overcome high-cost commercial boundaries within optical design have been of significant interest [4–10]. One such area of interest for both high-end and low-cost developments is super-resolution optical microscopy, with continuous developments in imaging beyond the diffraction limit both deeper and faster underlining this area of research. Among the wide variety of techniques developed, Image Scanning Microscopy (ISM) emerged to combine the principles of confocal microscopy with a pixel reassignment methodology to achieve resolution improvements beyond the conventional confocal microscope, achieving a 4-fold improvement to the signal-to-noise ratio (SNR) [11,12]. By using 1D or 2D laser-scanning approaches in fluorescence imaging and re-assigning the captured signals to account for parallax using a detector array, ISM can improve the lateral resolution of the system by up to a factor of √2 [13,14]. Image scanning microscopy has since been developed both commercially with low numbers of fast detector arrays, e.g. the Zeiss Airyscan detector [15], and in research settings to yield higher temporal resolution and sensitivity to detected photons, primarily through the incorporation of single photon avalanche detector (SPAD) arrays [16], incorporating also quantum mechanical detection methods [17]. With the inclusion of image deconvolution methods, the ISM method produces a doubling of the lateral imaging resolution of the microscope.

A key development in ISM, introduced by York *et al*. [18], made use of a microlens array – an optic which itself is made of multiple lenses smaller than the total optic diameter - to create multifocal structured illumination microscopy (mSIM) as an approach to improve the temporal resolution of the ISM technique. The microlens array is used to generate multiple parallel focused excitation points which are scanned across the imaging field of view, providing full field of view imaging speeds of around 1 Hz. The imaging speed has since been further improved using two similar microlens arrays for excitation and detection respectively to develop an all-optical method termed instant SIM (iSIM) [19], achieving super-resolution 2D imaging speeds surpassing 100 Hz. As the most affordable approach for multifocal illumination generation, commercially available microlens arrays are typically manufactured using high-precision lithography [20,21], and as such they offer excellent optical performance, though customisation of the array geometry is only possible at significant economic cost to the end-user. Lower cost commercial microlens arrays are available, though at the expense of optical excitation efficiency where lenslet diameters on the millimetre scale increase the number of required images and therefore decrease the temporal resolution. However, with increased lenslet spacing (pitch) the optical sectioning capabilities of the technique improve compared to a tighter lenslet pitch. As such, more efficient designs are possible, which can maximise the lenslet pitch while maintaining high optical sectioning capabilities.

Additive manufacturing approaches have so far shown to be capable of reducing costs in optical microscopy and providing user-specific custom designs [22–25], largely through 3D printing of optomechanical components [26–28]. One previous barrier to 3D printing high-quality optics was the required use of pulsed lasers for two-photon polymerisation [29,30]. More recently however, 3D printed optics have shown promise in optical imaging with the unique benefit of application-specific, customisable and freeform lens design [6,31,32] while maintaining low costs. Using commercially available, low-cost desktop stereolithography (SLA) printers and consumer-grade vat polymerisation printers (VPP), 3D printed optics have so far shown success in manufacturing intraocular contact lenses to correct for colour blindness [33], as well as in microscopy applications [34] and as imaging objectives for brightfield and fluorescence biology [10]. They have also been extensively characterised in terms of their material and optical properties [6,35–37]. Non-imaging optics within optical microscopy have also been produced using desktop printer modifications [38,39]. However, desktop 3D printed optics are yet to show their application within biological fluorescence excitation for super-resolution optical microscopy.

As additive manufacturing approaches for imaging-performance optics are well suited to provide lower cost custom optical elements, we present the design and manufacture of a custom 3D printed hexagonal (or ‘honeycomb’) lenslet array geometry and the integration of the optic into a custom small footprint (< 60 cm x 60 cm) mSIM system for fluorescence imaging. This custom mSIM microscope integrates Microelectromechanical Systems (MEMS) technology for array beam-scanning, and facilitates the evaluation of three different lenslet array designs: a high-end commercial microlens array (ML-1) with 250 µm lenslets and focal length (f) = 7.5 mm; a budget commercial lenslet array with 1.2 mm x 1.6 mm size lenslets (ML-2) and f = 4.7 mm; and the custom designed 3D printed honeycomb lenslet array (ML-3) with 1.2 mm pitch lenslets and f = 5 mm, with the lenslets configured in the honeycomb pattern to decrease the space between focal spots and therefore improve the temporal resolution compared to the more widely available commercial square or rectangular array designs. We evaluated the potential of additive manufacturing for lenslet array fabrication by conducting a comparative analysis of commercial and lab-fabricated optics and their respective impact on the performance of mSIM as a resolution-doubling microscopy method.

## 2. Materials and Methods

### 2.1 3 D printing and post-processing protocol

Manufacturing of optical quality lenslet arrays used 3D printing and silicone moulding. Traditionally, moulding uses a high-quality master copy to obtain precise moulds mimicking the surface roughness and shape of the master copy. This also works well as a technique for optical quality additive manufacturing, though the requirement of a master copy slightly reduces the economic and customisation benefits of 3D printing unless numerous copies of that specific optic are required. Nevertheless, 3D printing and spin-coating creates high quality surface roughness parts that can be used for custom lens designs. The custom design honeycomb array was designed in AutoCAD Inventor 2024 software and exported as a.STL file. The design files were translated to printer-readable file formats using a free slicer software (Chitubox basic) and printed using a consumer grade VPP 3D printer with 22 µm pixel size (Phrozen Sonic Mini 8K S) and 10 µm layer step size. The master copy was manufactured using a low-cost commercial resin (Anycubic, High Clear) and printed at a 90° angle to the print-bed to take full advantage of anti-aliasing features within the slicing software.

The fabrication process to obtain optical quality non-planar components, and specifically here the master copy from which we mould, is shown in Figure 1. The initial 3D printing step in A produces a print with the ‘staircase effect’ with layers equal to the chosen axial resolution. It is followed by a cleaning step B where the completed print was cleaned with 100% isopropyl alcohol (IPA) for a maximum of 10 minutes and blow-dried with compressed nitrogen. In step C the curved surface was spin-coated with a secondary clear resin (UV resin “Crystal clear”,Vida Rosa) using a friction-only affordable (∼£2000) spin-coater (Ossila L2001A3-E463-UK) at 6000 RPM for 10 seconds. The secondary resin is used to minimise curing time while maintaining high surface roughness quality [6,10]. The coated master mould was outgassed as shown in step E using a 55°C oven for 48 hours. Once the 3D print has been outgassed, the master copy is ready to be used to create the master mould with a low-cost commercially available two-part silicone mixture. The mixture was poured over the master copy as per step F and left to set for at least 24 hours at room temperature. The result is a flexible mould with inverse optical geometry to the master copy, matching the optical-quality surfaces of the original print. The master mould was subsequently filled with Vida Rosa resin as shown in step G, with air bubbles either pipetted away or left to pop over time before UV curing.

**Figure 1.**
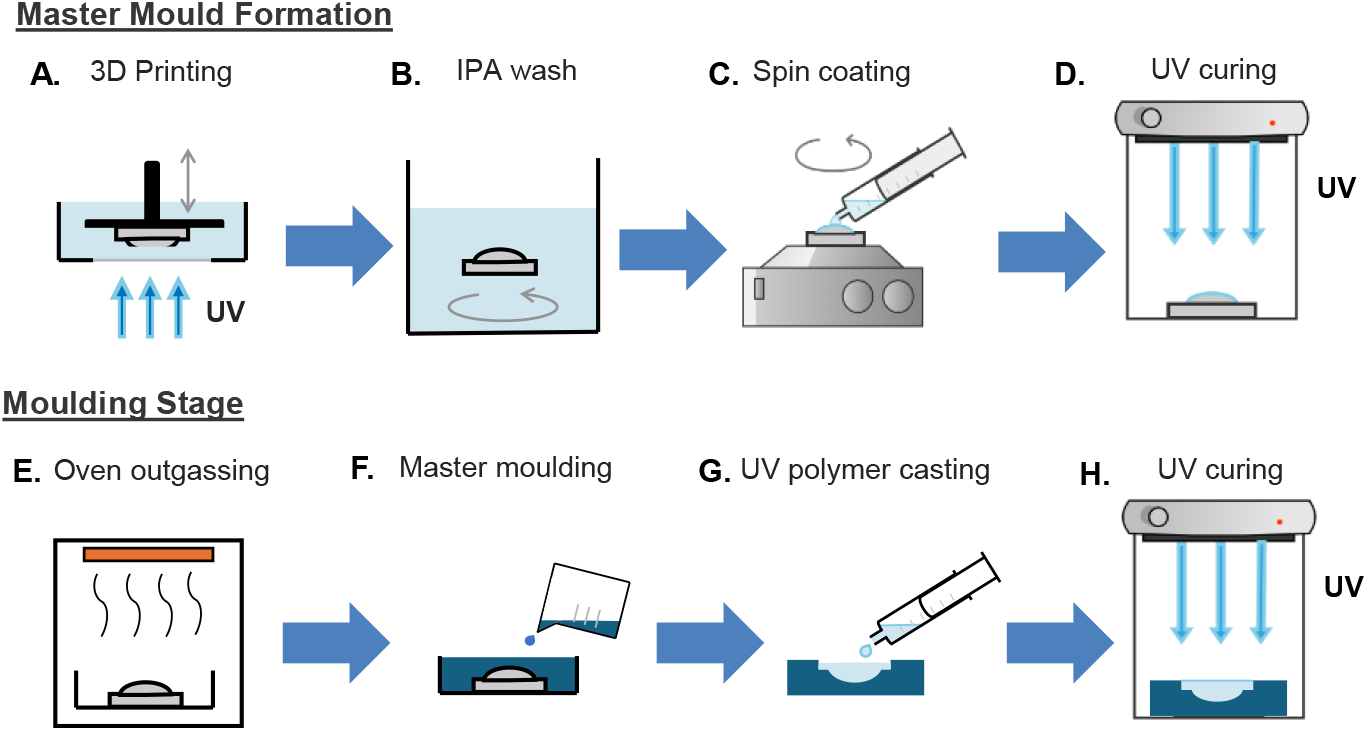
Schematic of 3D printed optics two-stage manufacturing process. (A)-(D) Manufacturing a master copy with optical quality non-planar surface; (E)-(G) Manufacturing a moulded optic using the master copy; (A) 3D printing in a layer-by-layer technique using incident UV light; (B) Cleaning stage using isopropyl alcohol (IPA) to remove residual resin; (C) spin-coating non-planar surface by pipetting resin onto surface prior to spinning; (D) non-planar spin-coated surface cured with UV light; (E) Oven outgassing of the master copy; (F) Silicone moulding to create a master mould; (G) UV curing resin casting from the master copy; (H) UV curing to complete the optically clear moulded lens.

To ensure the planar side of the optic had suitable surface roughness to act as an optic, a Vida Rosa spin-coated glass slide was gently applied to this surface, with the spin-coating step used to minimise air bubble generation between the slide and resin within the mould. The mould was then cured for 30 minutes, shown in stage H, to ensure complete curing throughout the bulk optic and to the edges of the mould. Once curing is complete, the moulded honeycomb lens array can be easily removed from the silicone and the glass slide removed through manual leveraging.

To minimise scattering at the boundaries of each lenslet, an opaque pinhole array was designed in the CAD software with matching geometry to the 3D printed honeycomb array. This was then 3D printed using ABS-like Black Resin to act as optical blocking layer and secured onto the non-planar lenslet surface.

To obtain surface roughness measurements for each of the lenslet arrays, a Tencor Alpha Step IQ Stylus profiler with a 5 µm tip measured a parametric sweep across each optic. A previously documented MATLAB script [10] was used to evaluate radius of curvature mismatches between optics and a theoretical ideal spherical curve. The contact approach was limited to measure one lenslet at a time when measuring the mm-scale lenslet arrays due to the limited axial measurement range while maintaining a high accuracy axial measurement resolution.

### 2.2 3D Printed mSIM processing pipeline

To obtain the mSIM processed image datasets, an image processing pipeline was made in Python 3.8.19 as shown in the schematic in Figure 2. The script processes a diffraction-limited laser-scanned image stack into an image stack featuring the pixel reassignment benefits of mSIM. The laser-scanned image stack itself is also combined into a diffraction-limited maximum intensity projection image for comparison against the mSIM processed result. The script runs on the assumption that a benchmark image calibration stack from a homogenously fluorescent slide is loaded first, though this step can be skipped by using the sample image stack in the absence of system data from a homogenous fluorescent slide. The pipeline then determines maximum intensity coordinates within each image, obtaining the centre point of the focused illumination originating from each lenslet element. Each maxima detected in a 2D image is separated by a user-defined pixel distance based on the lenslet spacing and resultant excitation sparsity, reducing the possibility of interference from hot pixels. With each intensity maximum located in the benchmark image stack, the sample image stack is then processed similarly, though with the option of including a maxima refiner; in essence a smaller ROI that the maximum intensity location will be searched for based around the maxima found in the benchmark stack. This can help to account for lateral stage drift. Each sample fluorescence maxima are then all cropped to a predefined ROI and convolved with a Gaussian blur with σ = 1.5 in the Python script. An additional intensity flat-field correction across the lenslet array elements is incorporated, normalising the intensities to a maximum value, to reduce intensity inhomogeneities scanned across the sample. With each ROI processed, the cropped fluorescent regions are reduced by a factor of 2 by sequentially adding them onto an ‘empty’ image with double the dimensions of the original benchmark or sample image stacks, creating the pixel reassignment process. By combining each resulting image into a single maximum intensity projection, the final mSIM 2D dataset is generated [13,40]. To obtain the optional full 2-fold resolution improvement, a Richardson-Lucy deconvolution step was used in a separate MATLAB script. The script used 5 iterations of the deconvolution algorithm with an experimentally measured point-spread function as basis, which was determined through the image of a laser-scanned sub-resolution bead.

**Figure 2.**
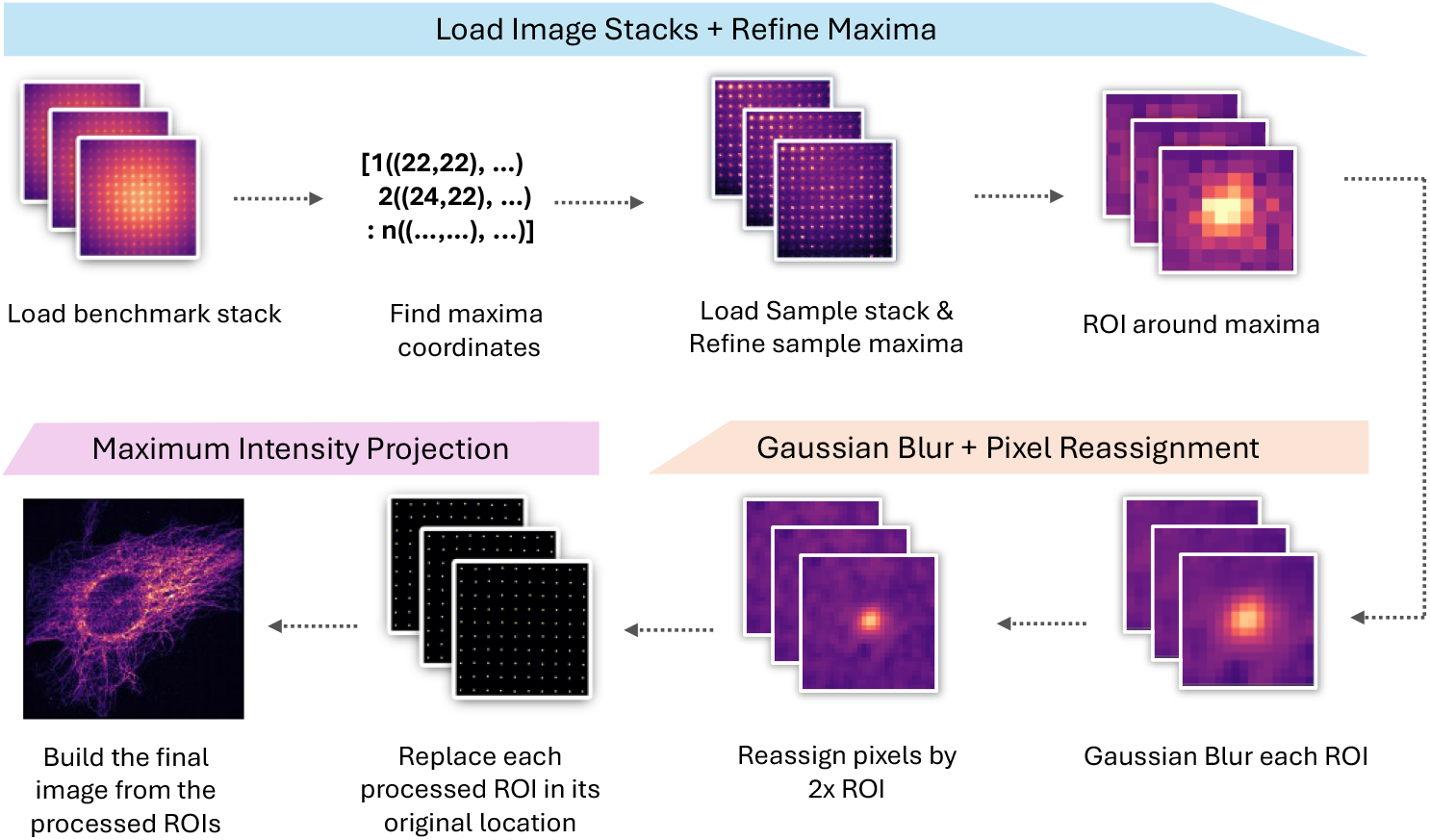
Image processing pipeline for mSIM image improvement. A benchmark stack is used to obtain reference maximum intensity coordinates. The coordinates are used to refine the maximum intensity coordinates from the sample image. Regions of interest (ROI) are cropped from the sample image and convolved using a Gaussian filter. The ROI is reassigned by upscaling with a factor of 2 and replaces the original unprocessed ROI at its original location. A maximum intensity projection results in the final processed image.

### 2.3 mSIM setup for custom optic integration

The constructed custom mSIM system (Figure 3) is built around a low-cost 488 nm laser diode excitation source (Odicforce OFL299-1), a 60x 1.42 NA oil immersion objective (Olympus PlanApo) and an industrial CMOS camera (IDS U3-3060CP). The single-mode fibre-coupled excitation laser is collimated using an air-spaced doublet collimation package (Thorlabs F810FC-543) and the resulting beam expanded by a 1.25x telescope (Thorlabs AC254-060-A and AC254-075-A) to fully fill the lenslet array under test. The lenslet array is mounted in a rotation plus tip-tilt mount (Edmund Optics 36-636) combined with a x-y-z translation stage (Thorlabs DT12XYZ/M) to be able to adjust for the varying focal lengths between different microlens implementations. The two commercial microlens arrays are the high-end 250 µm pitch array (Viavi MLA-S250-f30) and the budget commercial 1.2 mm by 1.6 mm array (Thorlabs MLA1). A pair of relay lenses (Thorlabs AC254-030-A) after the microlens array allow placement of an additional aperture that ensured that the excitation from each lenslet was both circular in profile and fit onto the 2 mm diameter MEMS micromirror used for 2D beam scanning. The MEMS micromirror allowed rapid (5 ms step response) yet precise (∼ 4 nm step size) positioning at the sample plane. Further 1” achromatic lenses are used to additionally adjust the beam size to fit the MEMS scanner aperture, with a final telescope (Thorlabs AC254-030-A and AC254-150-A) used to overfill the back-aperture of the 60x excitation objective. The IDS cockpit software was used to capture images at each scan location, combined with a custom Arduino program to control the MEMS scanner position angle and rising edge camera trigger. The microlens arrays used for comparing the 3D printed approach to commercial counterparts are introduced into the system via a flip mounted tip-tilt mirror on a 3-axis stage. This allows for the microlens arrays to be translated laterally and axially and supports fast swapping between widefield and laser-scanned imaging modalities.

**Figure 3.**
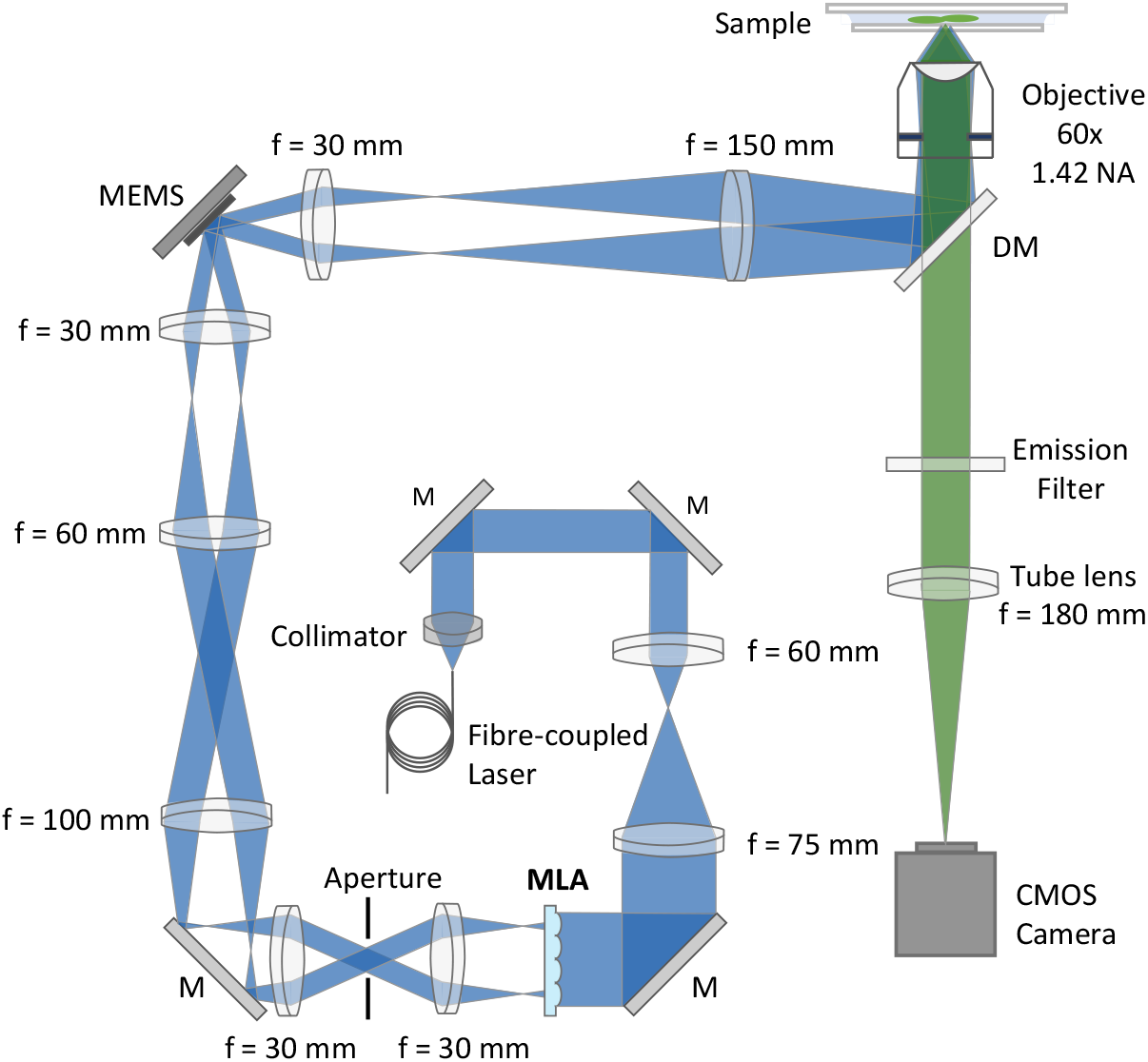
Schematic of the constructed mSIM microscope featuring a fibre-coupled 488 nm laser, microlens array, MEMS scanner, 60x 1.42 oil immersion objective, and CMOS camera. MLA – microlens array; M – Mirror; MEMS – microelectromechanical systems; DM – Dichroic mirror.

### 2.4 Sample preparation

For point spread function evaluation a sample of 505/515 excitation/emission yellow-green 175 nm nanobeads (PS-Speck™ Microscope Point Source Kit) was used. A 1:5 bead suspension was prepared using deionized water, and a 20 µL droplet of the suspension was placed on a #1.5 coverslip before being air-dried. ProLong Glass antifade mountant (Invitrogen P36980), with a refractive index of n=1.52, was applied to the beads after 2 hours of air-drying, with the coverslip then pressed onto a glass microscope slide and left to set for 24 hours. After setting, the coverslip was sealed on the microscope slide with clear nail varnish.

A commercial fixed bovine pulmonary artery endothelial (BPAE) cell slide (Invitrogen FluoCells Prepared Slide #1 F36924) was imaged with the custom microscope to validate the mSIM processing results. The slide was excited through a dichroic mirror (Chroma ZT405/488/561/640rpcv2) at 488 nm with emission at 520 nm and a band-pass emission filter in the detection path (Thorlabs FBH520-40) with 40 nm bandwidth. For the focused excitation positioning benchmark datasets required for mSIM processing, a uniform fluorescent slide was used (Thorlabs FSK2).

## 3. Results

The custom optic design for integration into the mSIM microscope is shown in Figure 4(a) and compared to a schematic of the low-cost commercial microlens array. The 3D printed array features a honeycomb geometry to optimise the functional area over a 1” diameter, opposed to the more regularly available commercial rectangular array design. This is shown in more detail in Figure 4(b), highlighting that the rectangular array features larger areas of unfilled space in comparison to the custom honeycomb configuration of lenslets. Therefore, the centre-centre distance between lenslets is reduced, also reducing the scan time for a 2D laser-scanned image. Due to the addition of a 3D printed array mask with 1 mm lenslet clear aperture the effective fill-factor of ML-3 is reduced to ∼83%, calculated as the effective optical area as a function of lenslet pitch. While this reduced fill-factor effects the overall light-throughput, it enhances contrast as it minimised scattering across the lenslet boundaries. The equivalent fill factor for ML-1 and ML-2 are ∼52% and ∼80% using the same calculation and assuming a similar proportion of lenslet mask across each array. Figure 4(c) highlights the spin-coated 3D printed optic master copy as well as the resultant silicone mould and final fabricated microlens array. A closer inspection of example lenslets for each microlens array is presented in Figure 4(d)-(f) with the high-end commercial ML-1 exhibiting sub-50 nm mismatch from an ideal spherical fit. In contrast, the lower cost ML-2 and the 3D printed honeycomb ML-3 both exhibit similar spherical profiles, with mismatches to the spherical fit generally below 1 µm. The nominal manufacturer quoted radius of curvature for the 250 µm microlens array could not be found, though the designed values for the ML-2 and the designed 3D printed honeycomb array are 2.7 mm and 3 mm respectively. The measured radius of curvature extracted using a spherical fitting function were 3.46 ± 0.05 mm, 2.63 ± 0.06 mm, and 2.46 ± 0.13 mm from the commercial 250 µm pitch ML-1, the commercial ML-2, and the 3D printed honeycomb ML-3 respectively after measuring three lenslets in each array chosen randomly across the optic surface.

**Figure 4.**
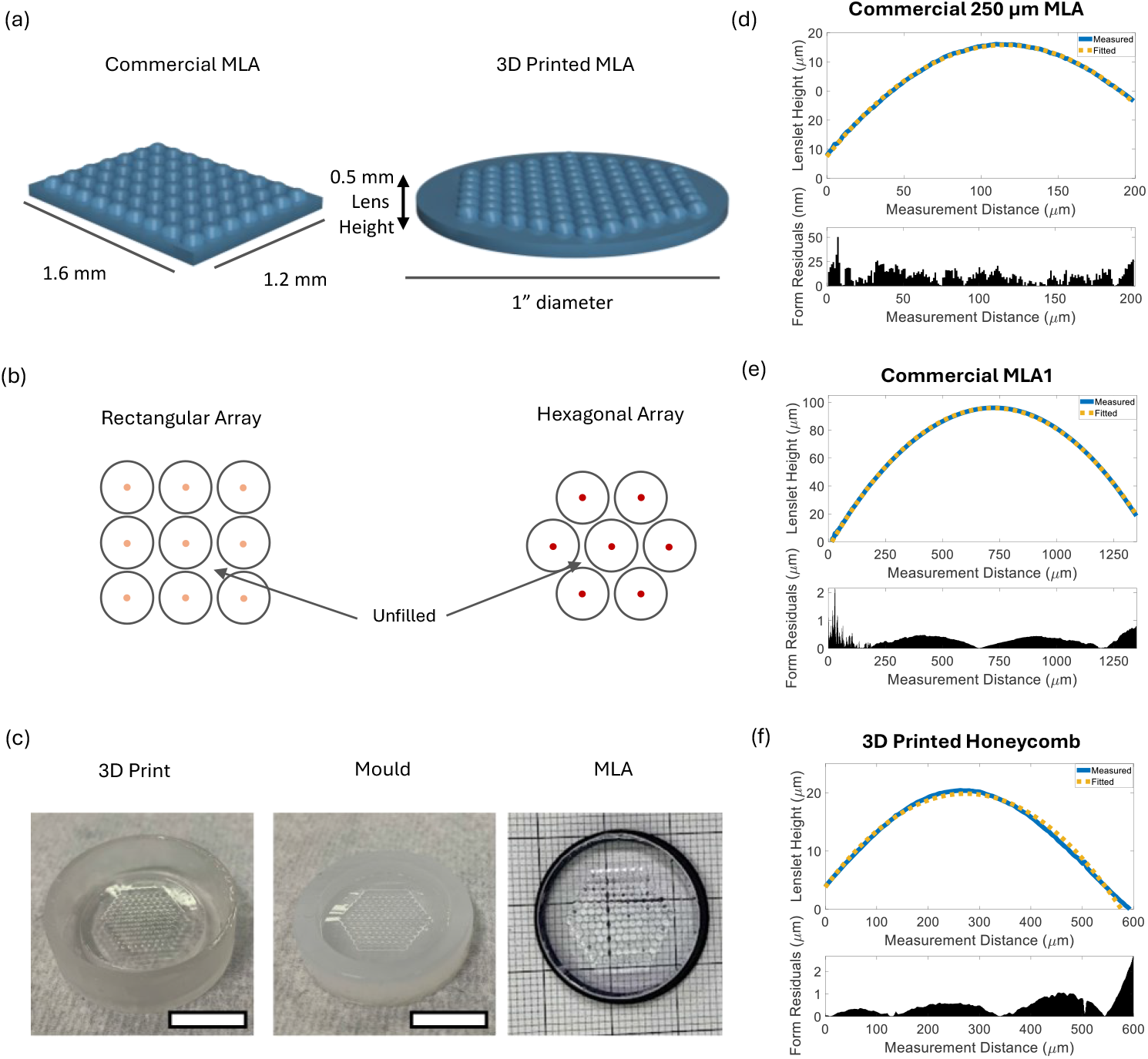
Microlens array schematic showing the design differences between the low-cost commercial array and the 3D printed array. (a) CAD design overview of the low-cost commercial rectangular lenslet array and the custom honeycomb geometry lenslet array design; (b) Schematic showing the relative density of lenslets in the rectangular and hexagonal arrays as a factor of unfilled area; (c) Photographs of the 3D printed master copy which has been coated for optical clarity, the master mould made from the master copy using consumer-grade silicone, and the resulting microlens array after using the silicone mould for manufacturing. Scale bars = 15 mm.; (d) – (f) surface profiles and spherical fits for the commercial 250 µm ML-1, commercial ML-2 and 3D printed ML-3 including the residual between the measured profile and a spherical fit.

To quantify the resolution of the designed mSIM system, the 175 nm diameter bead specimen was imaged using the widefield and laser-scanning modalities for each microlens array configuration. This provides lateral resolution information to evaluate how microlens array spacing influences the performance of mSIM as an optical resolution enhancement method. Images were captured at 5 ms exposure time with analogue and digital gain values of 10 and 1 respectively within the IDS cockpit software. In widefield, the 175 nm beads shown in Figure 5 exhibit an average full width half maximum (FWHM) value between ∼229 nm ± 11 nm and 242 nm ± 11 nm, showing similar performance within the error margin across all tests. Like the widefield results, the diffraction-limited laser-scanned maximum intensity projection in Figure 5(a) features an average bead FWHM of 235 nm ± 7 nm for the 250 µm ML-1 without deconvolution. The lower cost commercial ML-2 laser-scanned results show a similar FWHM of 247 nm ± 13 nm and the 3D printed honeycomb ML-3 shows a similar average bead FWHM of 246 nm ± 7 nm in the laser-scan modality both without deconvolution. When processing the scanned fluorescence signals using the developed mSIM Python script, average resolution improvements within the anticipated 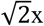 range were measured from the bead PSF FWHM of 166 nm ± 7 nm for the commercial 250 µm ML-1 in Figure 5(a), when measured across 5 individual beads chosen at random across the field of view and without the addition of deconvolution. When using the low-cost commercial ML-2 shown in Figure 5(b) the mSIM processed beads have an average FWHM value of 186 nm ± 6 nm when measured across 5 individual beads chosen at random across the field of view without deconvolution. Similarly, in the honeycomb geometry in Figure 5(c), the average bead FWHM in the mSIM dataset is 178 nm ± 9 nm when measured across 5 individual beads chosen at random across the field of view also without deconvolving the bead data. The bead FWHM results match the theoretical widefield diffraction limit of 224 nm, and the processed results also closely match their theoretical maximum resolution improvement of 174 nm. Additionally, the line profiles for each of the processed and unprocessed microlens array datasets shown in Figure 5 further exemplify mSIM processing improvements using the commercial and printed optics, highlighted through the narrowing of bead PSFs compared to the widefield and laser-scanned data.

**Figure 5.**
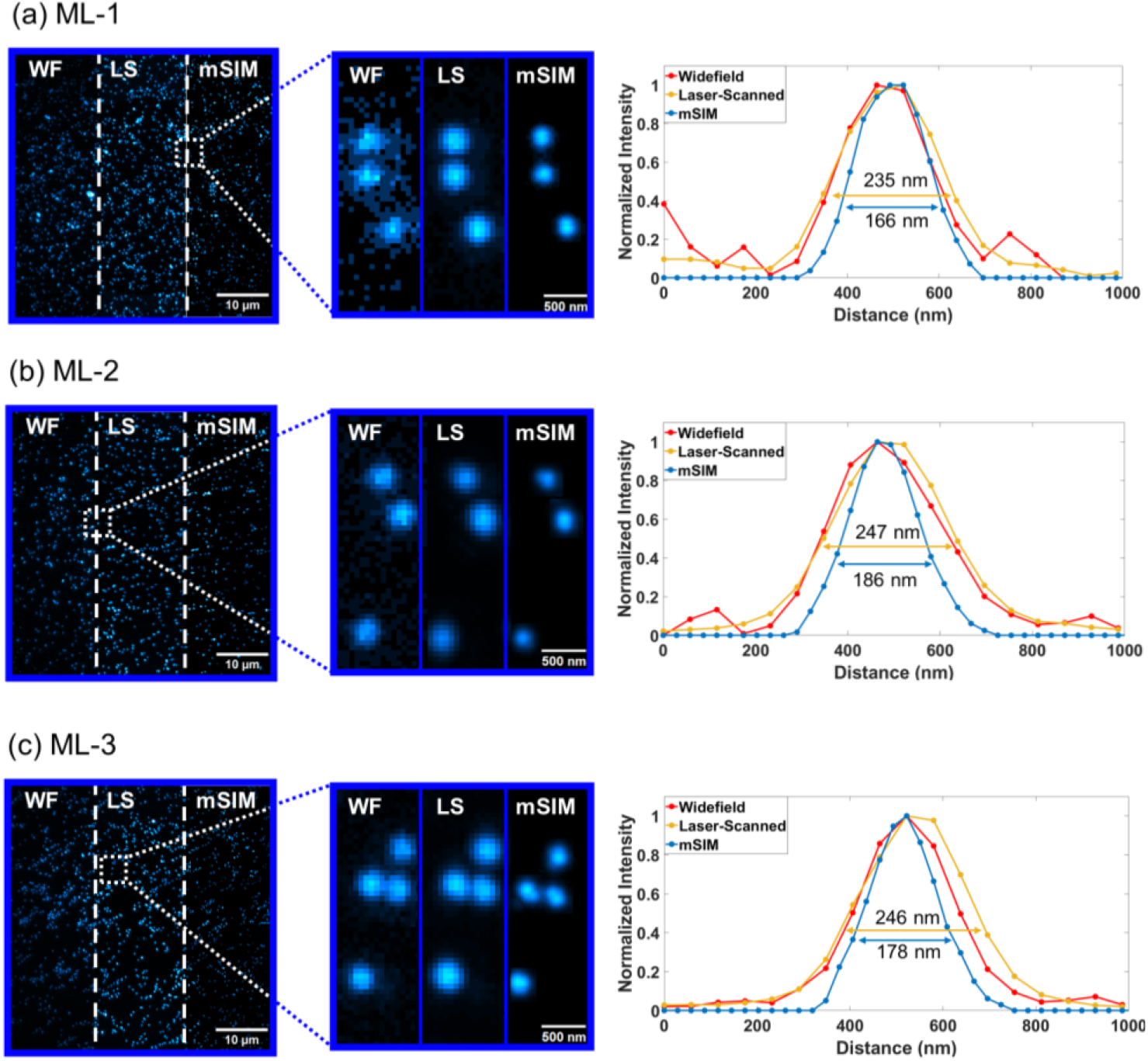
Diffraction-limited 175 nm fluorescent beads imaged via widefield (WF), laser-scanning (LS) and mSIM modalities. (a) 175 nm beads imaged using the commercial 250 µm ML-1 with full FOV, digital ROI and corresponding bead line-plot and FWHM; (b) 175 nm beads imaged using the low-cost commercial ML-2 with full FOV, digital ROI and corresponding bead line-plot and FWHM; (c) 175 nm beads imaged using the 3D printed honeycomb ML-3 with full FOV, digital ROI and corresponding bead line-plot and FWHM.

The image performance of the mSIM system using 3D printed optics was also characterised using fixed cell samples from a commercial fixed BPAE specimen, with α-Tubulin labelled using AlexaFluor 488 antibody. Images of the BPAE cells were captured using 5 ms exposure time and analogue and digital gain values of 20 and 5, respectively, using the IDS cockpit software. Tubulin filament thickness and signal-to-background ratio (SBR) were measured across five areas in the BPAE images at identical locations for the deconvolved laser-scanned and deconvolved mSIM results and averaged. Figure 6 shows the widefield, laser-scanned and mSIM images each under the same imaging conditions and power for the three microlens array types used. The images have been contrast enhanced in FIJI exclusively to enhance visual comparison. Figure 6(a) shows superior background rejection to the widefield results when using the 250 µm ML-1 array, resulting in higher contrast between the imaged tubulin and the background. Within the digitally zoomed ROI especially, the mSIM processed tubulin signal-to-background and resolution show additional notable improvements to the laser-scanned results, supported by the corresponding intensity cross-sections. Using the ML-1 array the average improvements to signal-to-background ratio (SBR) from laser-scanned to mSIM are 2.57x, with tubulin filament width resolved to 137 nm ± 11 nm when compared to 199 nm ± 21 nm without mSIM processing. Using the ML-2 array, comparable improvements in signal-to-background and resolution are observable, however, between the two commercial microlens arrays, the background rejection is improved in the sparser lenslet array dataset of ML-2. Specifically, ML-2 shows an average SBR improvement of 3.74x compared to the deconvolved laser-scanned results, with an average tubulin filament width of 133 nm ± 11 nm measured in comparison to the tubulin without mSIM processing which has 178 nm ± 8 nm filament width. The 3D printed honeycomb ML-3 data is shown in Figure 6(c) and the corresponding line profile shows close similarities in spatial resolution and contrast improvements to the results from both commercial optics when imaging and processing the BPAE sample. The ML-3 optic shows average SBR improvements to the laser-scanned results of 3.78x, with the average tubulin width resolving to 134 nm ± 9 nm from the deconvolved laser-scanned results of 180 nm ± 12 nm. Additionally, the normalised intensity differences in comparison to the laser-scanned results in all three line profiles combined with the distinguishable difference in lateral details shown by increased peak-trough sections both exemplify the similarities in resolution and contrast improvements from all three optics.

**Figure 6.**
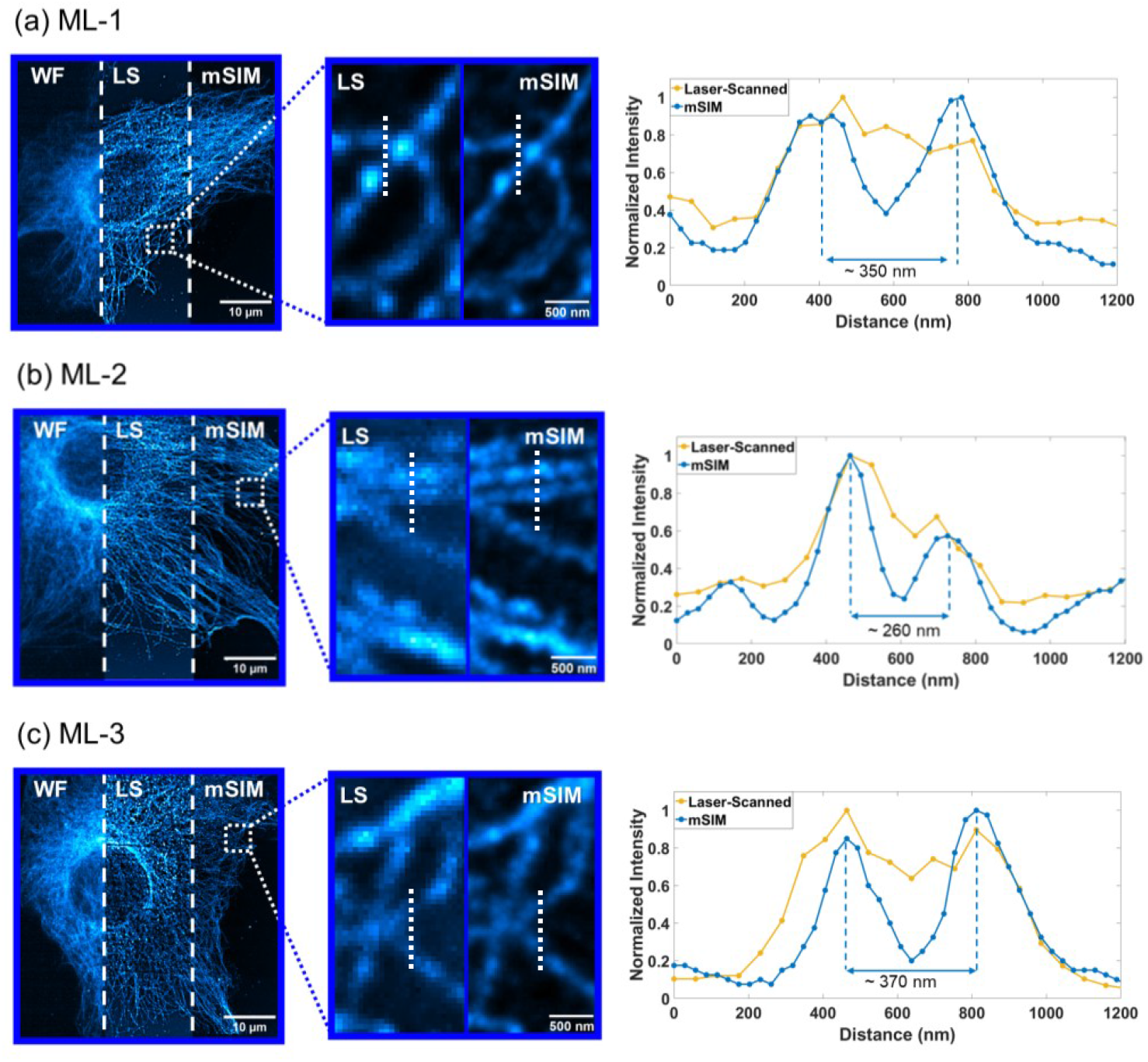
Images of fixed BPAE cells with α-Tubulin labelled Alexa Fluor 488 imaged via widefield (WF), laser-scanning (LS) and mSIM modailities. (a) BPAE imaged using the commercial 250 µm ML-1 array with full FOV, digital ROI and corresponding tubulin line-plot; (b) BPAE imaged using the commercial ML-2 array with full FOV, digital ROI and corresponding tubulin line-plot (c) BPAE imaged using the 3D printed ML-3 array with full FOV, digital ROI and corresponding tubulin line-plot.

Shown in Figure 7 are the respective illumination inhomogeneities after maximum intensity projection for the 250 µm ML-1 (a), ML-2 (b) and 3D printed honeycomb ML-3 (c). There is minimal inhomogeneity in the illumination across the selected ROI of the BPAE image when using the commercial 250 µm ML-1. In contrast to this, more significant intensity variations can be seen between the arrow locations in the ROI in Figure 7(b) and (c) resulting from the inhomogeneous illumination scanned across the fluorescent sample, despite some computational flat-field correction. Additionally, some striping can be observed in the full field of view images of Figure 7(b) and (c) in particular, primarily due to the lack of autofocus correction as the longer scan times contributed to some axial drift.

**Figure 7.**
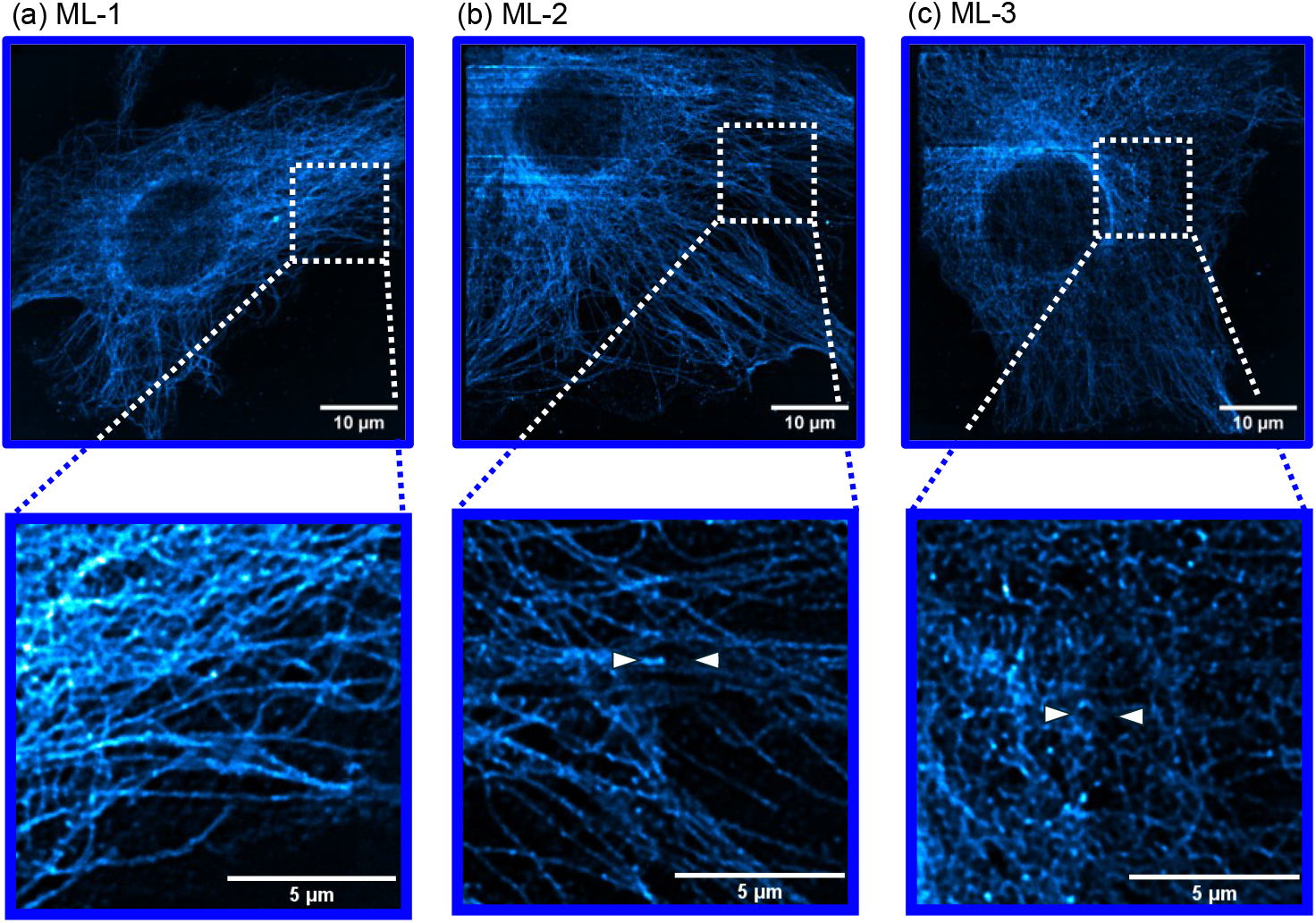
Illumination inhomogeneities across all three microlens arrays when imaging fluorescent BPAE. (a) – Intensity variations resultant from the commercial 250 µm ML-1 with digitally-zoomed ROI; (b) - Intensity variations resultant from the commercial ML-2 with digitally-zoomed ROI; (c) - Intensity variations resultant from the 3D Printed honeycomb ML-3 with digitally-zoomed ROI.

## 4. Discussion

The surface profiles from the lenslet arrays in Figure 4(d)-(f) show that the 3D printed honeycomb lenslets as well as the commercial variants are well matched to ideal spherical fits. However, at the border between lenslet elements the 3D printed honeycomb lenslet array features a smooth transition between elements, slightly reducing the clear aperture of each lenslet. The smoothing at the lenslet edges are an added reason for the addition of the honeycomb geometry-matched opaque pinhole array, as inhomogeneities from the clear resin spin-coating process at the locations between lenslets become present in the sample. By adding the 3D printed pinhole array component, the additional background and lenslet-edge sample illumination were removed. However, the surface profile data does suggest that the lenslet array can be manufactured to optical quality without the requirement of costly two-photon techniques as shown previously [29,30], though instead at the expense of minimum-achievable lenslet pitch size. The radii of curvature for the ML-2 optic matches closely the nominal value, with the 3D printed honeycomb ML-3 outside of the design specification slightly more than in the commercial case. The radius of curvature for the 3D printed ML-3 lenslets is expectedly lower than the designed curvature likely due to excess thickness on the lenslet surface contributing to a shorter curvature. This can be optimised to be removed through iterative lenslet curvature redesign while maintaining all fabrication parameters, or by spin-coating the curved surface with a faster RPM to obtain a thinner layer of covering resin and therefore shallower radius of curvature. In this work the maximum spin-coating speed was limited to 6000 RPM and we could not therefore hone the optic curvature this way. When fabricating the optic, volumes of resin < 20 ml were used with approximately 10 g of the total silicone mixture required; pricing the resulting optic relative to these material costs therefore makes the final manufactured lenslet array ∼ £0.50. However, as this is the cost to manufacture the master copy, master mould and final optic, the price of each optic manufactured decreases as the mould can be used repeatably. To achieve a comparable signal-to-background when imaging samples using each array, the laser excitation power was adjusted for each microlens array optic as each microlens array absorbs and transmits the excitation light slightly differently due to differences in material and thickness. The results of the mSIM processing pipeline shown in Figures 5 and 6 exemplify the effectiveness of mSIM as a super-resolution method when used in conjunction with a low cost, 3D printed honeycomb lenslet array, as image quality improvements in SBR of ∼ 3.7x and resolution down to ∼ 134 nm are observed in fixed cell imaging and are comparable to results obtained with commercial microlens arrays. The mSIM processed datasets outperform the widefield and laser-scanned maximum intensity projection counterparts, matching closely the expected √2x resolution improvement in lateral resolution without deconvolution, and resolution doubling when combining mSIM processing with deconvolution, as also shown in conventional commercial-microlens array implementations of mSIM systems [18,19]. The widefield images in Figure 6 show higher background due to the identical imaging conditions to the laser-scanned images, which therefore contributes higher noise and scattering which significantly affects the resolution of the acquired image. By utilising laser-scanning and deconvolution methods with commercial and 3D printed honeycomb lenslet arrays, significant image quality improvements are observed, with the resolution and contrast enhanced beyond the optical diffraction limit through mSIM methods as expected and supported by the corresponding intensity plots in Figure 6. These results therefore show that biological imaging beyond the optical diffraction limit is possible using low-cost desktop 3D printed optics.

Figure 7 shows that for sparse lenslet arrays, excitation which is not uniform across the full illuminated FOV can result in inhomogeneous 2D image reconstructions in comparison to the smaller scan regions from the 250 µm ML-1. To correct for this, a flat top illumination profile would be beneficial. Additional reasons for the inhomogeneity observed across the BPAE sample, particularly for the 3D printed optic-case, may arise from inhomogeneities across the lenslets themselves originating from the fabrication process. Even with these potential inhomogeneities, microscope images of beads and cell slides beyond the optical diffraction limit have been shown using a 3D printed optical element at its core, enabling super-resolution microscopy approaches with affordable and customisable optical elements.

## 5. Conclusions

We have shown a custom 3D printed honeycomb lenslet array designed and manufactured using low-cost optical fabrication methods and used in super-resolution microscopy, specifically a custom mSIM setup. This allows for imaging with a resolution of 134 nm using a 60x 1.42NA objective, where mSIM processing and deconvolution provide the anticipated lateral resolution improvements of 2x with features beyond the optical diffraction limit of 246 nm resolved using the 3D printed optic. The use of custom designed, low-cost optics holds promise for future applications in alternative super-resolution microscopy methods as well as other areas of non-imaging optics. Therefore, it has been demonstrated that user-specific elements and illumination patterns are now accessible without sacrificing quality of results.

## Data availability

Data underlying the results presented in this paper are available in Ref 41

## References

1. J. W. P. Brown, A. Bauer, M. E. Polinkovsky, A. Bhumkar, D. J. B. Hunter, K. Gaus, E. Sierecki, and Y. Gambin, “Single-molecule detection on a portable 3D-printed microscope,” Nat Commun 10, 1–7 (2019).

2. T. Matsui and D. Fujiwara, “Optical sectioning robotic microscopy for everyone: the structured illumination microscope with the OpenFlexure stages,” Opt Express 30, 23208 (2022).

3. T. Liu, M. Rajadhyaksha, and D. L. Dickensheets, “MEMS-in-the-lens architecture for a miniature high-NA laser scanning microscope,” Light Sci Appl 8, (2019).

4. K. D. D. Willis, E. Brockmeyer, S. E. Hudson, and I. Poupyrev, “Printed Optics: 3D Printing of Embedded Optical Elements for Interactive Devices,” in Proceedings of the 25th Annual ACM Symposium on User Interface Software and Technology (UIST ‘12) (Association for Computing Machinery, 2012), pp. 589–598.

5. M. Elgarisi, V. Frumkin, O. Luria, and M. Bercovici, “Fabrication of freeform optical components by fluidic shaping,” Optica 8, 1501 (2021).

6. L. M. Rooney, J. Christopher, B. Watson, Y. S. Kumar, L. Copeland, L. D. Walker, S. Foylan, W. B. Amos, R. Bauer, and G. McConnell, “Printing, Characterizing, and Assessing Transparent 3D Printed Lenses for Optical Imaging,” Adv Mater Technol 9, (2024).

7. M. Hisham, A. E. Salih, and H. Butt, “3D Printing of Multimaterial Contact Lenses,” ACS Biomater. Sci.Eng. 2023, 9, 4381–4391 (2023).

8. Y. Xu, P. Huang, S. To, L.-M. Zhu, and Z. Zhu, “Low-Cost Volumetric 3D Printing of High-Precision Miniature Lenses in Seconds,” Adv. Optical Mater. 2022, 10, 2200488 (2022).

9. D. Gonzalez-Utrera, B. Villalobos-Mendoza, R. Diaz-Uribe, and D. Aguirre-Aguirre, “Modeling, fabrication, and metrology of 3D printed Alvarez lenses prototypes,” Opt Express 32, 3512 (2024).

10. J. Christopher, L. M. Rooney, M. Donnachie, D. Uttamchandani, G. McConnell, and R. Bauer, “Low-cost 3D printed lenses for brightfield and fluorescence microscopy,” Biomed Opt Express 15, 2224 (2024).

11. C. J. R. Sheppard, “Super-resolution in confocal imaging,” Optik (Jena) 80, 53–54 (1988).

12. S. Roth, C. Sheppard, K. Wicker, and R. Heintzmann, “Optical photon reassignment microscopy (OPRA),” Bioinformatics 2–7 (2012).

13. C. B. Müller and J. Enderlein, “Image scanning microscopy,” Phys Rev Lett 104, 1–4 (2010).

14. E. N. Ward and R. Pal, “Image scanning microscopy: an overview,” J Microsc 266, 221–228 (2017).

15. J. Huff, “The Airyscan detector from ZEISS: confocal imaging with improved signal-to-noise ratio and super-resolution,” Nat Methods 12, ii (2015).

16. A. Zunino, G. Garrè, E. Perego, S. Zappone, M. Donato, N. Vastenhouw, and G. Vicidomini, “Structured detection for simultaneous super-resolution and optical sectioning in laser scanning microscopy,” Nat Photonics 19, (2025).

17. U. Rossman, R. Tenne, O. Solomon, I. Kaplan-Ashiri, T. Dadosh, Y. C. Eldar, and D. Oron, “Rapid quantum image scanning microscopy by joint sparse reconstruction,” Optica 6, 1290 (2019).

18. A. G. et al. York, S. H. Parekh, D. Dalle Nogare, R. S. Fischer, K. Temprine, M. Mione, A. B. Chitnis, C. A. Combs, and H. Shroff, “Resolution doubling in live, multicellular organisms via multifocal structured illumination microscopy,” Articles Nature methods | 9, 749 (2012).

19. A. Curd, A. Cleasby, K. Makowska, A. York, H. Shroff, and M. Peckham, “Construction of an instant structured illumination microscope,” Methods 88, 37–47 (2015).

20. Y. Hu, Y. Chen, J. Ma, J. Li, W. Huang, and J. Chu, “High-efficiency fabrication of aspheric microlens arrays by holographic femtosecond laser-induced photopolymerization,” Appl Phys Lett 103, (2013).

21. S. Thiele, C. Pruss, A. M. Herkommer, and H. Giessen, “3D printed stacked diffractive microlenses,” Opt Express 27, 35621 (2019).

22. Q. Chen, J. D. Mangadlao, J. Wallat, A. De Leon, J. K. Pokorski, and R. C. Advincula, “3D printing biocompatible polyurethane/poly(lactic acid)/graphene oxide nanocomposites: Anisotropic properties,” ACS Appl Mater Interfaces 9, 4015–4023 (2017).

23. M. P. Chae, W. M. Rozen, P. G. McMenamin, M. W. Findlay, R. T. Spychal, and D. J. Hunter-Smith, “Emerging Applications of Bedside 3D Printing in Plastic Surgery,” Front Surg 2, 1–14 (2015).

24. F. Auricchio and S. Marconi, “3D printing: Clinical applications in orthopaedics and traumatology,” EFORT Open Rev 1, 121–127 (2016).

25. M. K. E. Cabello and J. E. De Guzman, “Utilization of accessible resources in the fabrication of an affordable, portable, high-resolution, 3D printed, digital microscope for Philippine diagnostic applications,” PLOS Global Public Health 3, (2023).

26. B. R. Patton, S. D. Grant, G. S. Cairns, and J. Wistuba, “Adapting the 3D-printed Openflexure microscope enables computational super-resolution imaging,” F1000Res 8, 1–13 (2019).

27. S. B. Tristan-Landin, A. M. Gonzalez-Suarez, R. J. Jimenez-Valdes, and J. L. Garcia-Cordero, “Facile assembly of an affordable miniature multicolor fluorescence microscope made of 3D-printed parts enables detection of single cells,” PLoS One 14, 1–17 (2019).

28. S. Amann, M. von Witzleben, and S. Breuer, “3D-printable portable open-source platform for low-cost lens-less holographic cellular imaging,” Sci Rep 9, 11260 (2019).

29. T. Gissibl, S. Thiele, A. Herkommer, and H. Giessen, “Two-photon direct laser writing of ultracompact multi-lens objectives,” Nat Photonics 10, 554–560 (2016).

30. L. Zhang, C. Wang, C. Zhang, Y. Xue, Z. Ye, L. Xu, Y. Hu, J. Li, J. Chu, and D. Wu, “High-Throughput Two-Photon 3D Printing Enabled by Holographic Multi-Foci High-Speed Scanning,” Nano Lett 24, 2671–2679 (2024).

31. B. I. G. A. Assefa, M. A. P. Pekkarinen, H. E. P. Partanen, J. O. B. Iskop, J. A. R. I. T. Urunen, and J. Y. S. Aarinen, “Imaging-quality 3D-printed centimeter-scale lens,” 27, 12630–12637 (2019).

32. L. D. Vallejo-Melgarejo, R. G. Reifenberger, B. A. Newell, C. A. Narváez-Tovar, and J. M. Garcia-Bravo, “Characterization of 3D-printed lenses and diffraction gratings made by DLP additive manufacturing,” Rapid Prototyp J 25, 1684–1694 (2019).

33. F. Alam, M. Ali, M. Elsherif, A. E. Salih, N. El-atab, and H. Butt, “3D printed intraocular lens for managing the color blindness,” Additive Manufacturing Letters 5, 100129 (2023).

34. J. Christopher, R. Craig, R. E. McHugh, A. J. Roe, R. Bauer, B. Patton, G. McConnell, and L. M. Rooney, “A 3D-printed optical microscope for low-cost histological imaging,” J Microsc 9 (2025).

35. S. Juodkazis, “Manufacturing: 3D printed micro-optics,” Nat Photonics 10, 499–501 (2016).

36. N. Vaidya and O. Solgaard, “3D printed optics with nanometer scale surface roughness,” Microsyst Nanoeng 4, (2018).

37. K. Sharma, T. Bora, and W. S. Mohammed, “Effect of solvent absorption on the optical properties of 3D printed methacrylate waveguide,” Opt Laser Technol 134, 106573 (2021).

38. G. Shao, R. Hai, and C. Sun, “3D Printing Customized Optical Lens in Minutes,” Adv Opt Mater 8, (2020).

39. X. Chen, W. Liu, B. Dong, J. Lee, H. O. T. Ware, H. F. Zhang, and C. Sun, “High-Speed 3D Printing of Millimeter-Size Customized Aspheric Imaging Lenses with Sub 7 nm Surface Roughness,” Advanced Materials 30, 1–8 (2018).

40. I. Gregor and J. Enderlein, “Image scanning microscopy,” Curr Opin Chem Biol 51, 74–83 (2019).

41. J. Christopher, “Data for Low-Cost 3D Printed Optics for Super-Resolution Multifocal Structured Illumination Microscopy”, Univ. of Strath. (2025) DOI: 10.15129/84502887-d283-4b05-bad6-2e9900bd3f57

